# Mean temperature determines whether winter variability accelerates or buffers energy loss

**DOI:** 10.64898/2026.03.11.711084

**Authors:** Jordan R. Glass, Sarah A. Waybright, D.M. Shayne Dodge, Ellen C. Keaveny, Sabrina A. White, Michael E. Dillon

**Affiliations:** Department of Zoology and Physiology, University of Wyoming, Laramie, WY, United States; Program in Ecology and Evolution, University of Wyoming, Laramie, WY, United States; Department of Neurobiology, Physiology, and Behavior, University of California, Davis, CA, United States; Department of Biology, University of North Carolina at Chapel Hill, Chapel Hill, NC, United States

**Keywords:** Diapause, resting metabolism, metabolic suppression, thermal sensitivity, *Bombus impatiens*, overwintering energetics, temperature variability

## Abstract

Winter survival in dormant animals depends on conserving finite energy reserves, yet winter temperatures fluctuate around shifting means. In ectotherms, metabolic rate increases exponentially with temperature, so thermal variability is expected to accelerate energy loss, with important consequences for overwinter survival and population persistence under climate change. However, it remains unclear whether dormant ectotherms can compensate physiologically for thermal variability. We overwintered *Bombus impatiens* queens under constant (2, 3, 4°C) or variable (2 ± 6°C or 4 ± 6°C) regimes for six weeks, then measured metabolic rates across a range of temperatures. The temperature dependence of metabolic rate shifted in response to thermal experience, but the direction of compensation depended on mean temperature: variability centered on 2°C elevated metabolic rate and increased thermal sensitivity relative to all other conditions, whereas variability centered on 4°C reduced metabolic rate and dampened thermal sensitivity relative to constant 4°C. We used these metabolic responses to simulate rates of lipid depletion and found that survival trajectories echoed physiological shifts: experiencing variability around 2°C would reduce subsequent survival time, whereas experiencing variability around 4°C would preserve subsequent survival even under variable future conditions. Thus, identical thermal variance produced opposite energetic outcomes depending on the mean temperature around which fluctuations occurred. Integrating both temperature means and variability is, therefore, essential for predicting overwintering survival in a changing world.

## Introduction

Temperature variability is a defining feature of natural environments. Daily highs and lows, seasonal cycles, and abrupt cold snaps or warm spells create dynamic thermal landscapes that strongly influence organismal physiology (1–7). For ectotherms, these fluctuations can be especially consequential because metabolic rate rises exponentially with temperature, such that repeated exposure to warmer temperatures disproportionately raises total energy expenditure (i.e., Jensen’s inequality or non-linear averaging; Denny, 2017; Jensen, 1906; Martin & Huey, 2008; Ruel & Ayres, 1999). As a result, fluctuating thermal regimes can increase mean metabolic rate relative to a constant regime at the same mean temperature (Fig. 1A), accelerating depletion of energy stores available for growth, reproduction, and survival.

**Figure 1.**
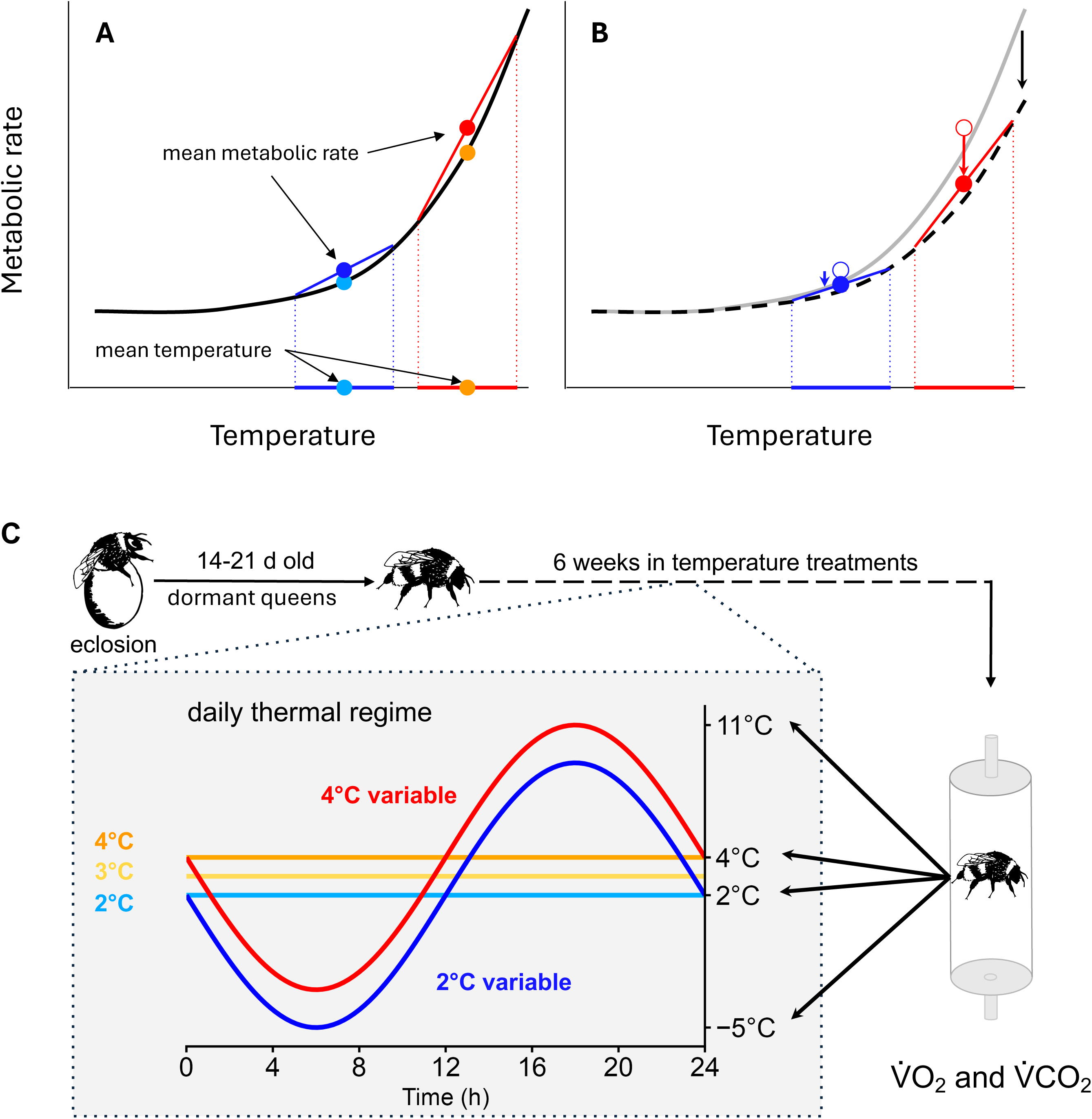
Energetic consequences of and compensatory responses to thermal variability. (A) The exponential relationship between metabolic rate and temperature results in higher mean metabolic rates (dark blue and red points) under variable temperatures (dark blue and red horizontal lines) than under constant temperatures (light blue and orange points). (B) Animals may respond through metabolic compensation, a shift in the metabolic rate-temperature relationship (dashed line), resulting in a reduction in mean metabolic rates (filled points) relative to previous values (open points). The magnitude of the compensation response and of the effect on metabolic rates may vary with mean temperature (red vs. blue points and lines). For both A and B, dotted vertical lines facilitate visualization of the translation of the range of temperature variability to the metabolic curve. (Redrawn after Denny, 2017). (C) Experimental approach to measuring compensatory responses to temperature variability. After eclosion, bumble queens were induced into dormancy before being placed in one of five treatments where they experienced daily thermal regimes that were constant (2, 3, 4°C) or variable (± 6°C at a mean of 2 or 4°C) for six weeks, at which point metabolic rates were measured at each of four temperatures.

However, some organisms can adjust their physiology in response to thermal variation through metabolic compensation, altering baseline metabolic demand and/or thermal sensitivity of metabolism in response to their thermal environment. Importantly, compensation is not unidirectional. In insects that remain active in cold environments, metabolic compensation may elevate metabolic capacity to sustain performance at low temperatures (12). During energetically constrained life stages, such as dormancy, compensation would be expected to reduce baseline metabolic demand or thermal sensitivity to conserve limited energy reserves (13). Although the energetic consequences of thermal variability are well documented in active insects – where fluctuations can reduce reproductive output (14, 15) – we know far less about whether dormant ectotherms can adjust their metabolism to buffer these costs, even though such responses could strongly influence overwinter survival and population persistence.

Dormant ectotherms must survive winter without feeding and, therefore, rely on finite energy reserves (16). These energetic constraints are further complicated by the fact that winter temperatures are rarely constant. Even within thermally buffered refuges such as soil burrows or hibernacula, individuals experience substantial variation, particularly during early and late winter (17–20). If dormancy physiology constrains metabolic adjustment, thermal variability should increase energetic expenditure through repeated exposure to warmer temperatures that partially reactivate metabolism and accelerate resource depletion (21).

Conversely, if dormant individuals retain the capacity for compensation, they may shift metabolic trajectories toward lower baseline demand or reduced thermal sensitivity, minimizing the energetic consequences of winter variability (19).

Crucially, the energetic consequences of variability likely depend not only on the magnitude of temperature fluctuations but also on the mean temperature around which variability occurs and how frequently temperatures cross physiologically important thresholds (22). Variability centered on relatively warm temperatures will increase cumulative energetic demand but may also provide sufficient time at permissive temperatures to enable compensatory remodeling or recovery (23, 24). In contrast, if variability is centered on colder temperatures, even brief warming events may repeatedly trigger metabolically costly physiological transitions without allowing enough time for compensation, generating cumulative costs that are not reflected in mean temperature alone (21, 25). Thus, thermal regimes with identical degrees of variability but different mean temperatures can yield strikingly different energetic outcomes, with consequences for overwinter survival and subsequent seasonal success.

Dormant bumble bee queens (*Bombus* spp.) provide a useful system for testing how thermal variability shapes metabolic regulation during winter dormancy. As solitary overwintering individuals, queens must conserve finite lipid reserves while avoiding freezing injury to survive winter, and overwinter survival directly determines spring colony establishment and population persistence (20). Understanding how mean temperature and thermal variability interact to shape overwinter energetics is therefore critical for predicting winter mortality under increasingly variable climates (21, 26).

To address this gap, we exposed overwintering *Bombus impatiens* queens to one of five ecologically relevant winter regimes (Fig. 1C): three constant cold treatments (2, 3, or 4°C) and two thermally variable treatments (2 ± 6°C or 4 ± 6°C). These treatments span temperatures commonly experienced by overwintering queens in soil refuges and encompass both typical diapause temperatures and periodic warming events observed during winter (20). This design allowed us to test not only whether thermal variability alters overwintering energetics, but also whether mean winter temperature alone induces metabolic compensation during dormancy. After six weeks of exposure, we measured metabolic rates (V[CO_2_ and V[O_2_) across four test temperatures (−5, 2, 4, and 11°C) to determine whether queens shift metabolic rate at a given temperature or alter the temperature sensitivity of metabolism, and whether these responses depend on mean temperature under both constant and variable thermal regimes.

We predicted that compensation would be strongest at higher mean temperatures, where queens should suppress metabolic demand to conserve finite energy reserves during diapause. We further predicted that variability would amplify this response, particularly at higher mean temperatures, where fluctuations more frequently reach temperatures at which metabolic rate rises steeply. Under this scenario, variability centered on 4°C should induce stronger metabolic suppression than variability centered on 2°C (Fig. 1). If queens compensate physiologically for thermal variability, these metabolic shifts should influence energetic trajectories during dormancy. We therefore expected individuals that reduce metabolic rate or thermal sensitivity to retain energy reserves longer and show slower lipid depletion than individuals unable to adjust.

## Results

### Thermal experience alters the temperature dependence of metabolic rate

Overwintering thermal regimes significantly altered the temperature dependence of metabolic rate in dormant bumble bees. Thermal regime (*P* = 0.003), measurement temperature (*P* < 0.001), and their interaction (*P* < 0.001) all significantly affected mass-specific CO_2_ production (spV[CO_2_) and O_2_ consumption (spV[CO_2_; *P* = 0.011, *P* < 0.001, *P* < 0.001, respectively; Table 1). Treatment effects were most pronounced at the warmest test temperature (11°C), indicating that early overwintering thermal history exerts its strongest influence on metabolic rate at higher measurement temperatures (Fig. 2A,B; Table 1).

**Figure 2.**
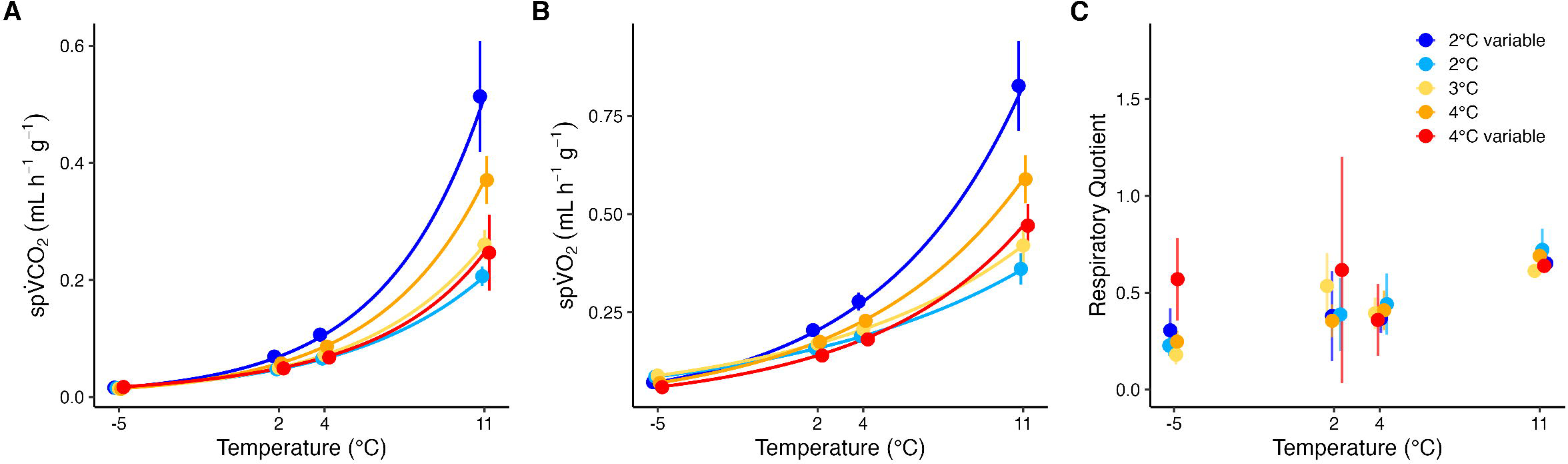
Early thermal experience alters the temperature dependence of metabolism in overwintering queen bumble bees. (A) Mass-specific CO_2_ production, (B) mass-specific O_2_ consumption (B), and (C) respiratory quotient (RQ = VCO_2_:VO_2_) measured at four test temperatures (−5, 2, 4, and 11°C) for queen bumble bees overwintered for six weeks under one of five thermal regimes (colors as in Fig. 1). For metabolic rates (A, B), points represent mean predicted values at each test temperature, averaged across bees (*n* = 18-22) within each treatment, with error bars indicating ± SE across bees; lines show mean Arrhenius relationships generated from per-bee nonlinear fits and averaged within treatments. For RQ (C), points show treatment-level estimated means with 95% confidence intervals.

**Table 1.** Rates of CO_2_ production (spV[CO_2_), O_2_ consumption (spV[O_2_), both mass-specific, and respiratory quotient (RQ) measured at 11°C, together with Arrhenius activation energy (*E*_a_) and intercept (ln *b*_0_) derived from individual-level Arrhenius fits for spV[CO_2_ and spV[O_2_. Values are estimated marginal means ± SE at 11°C for spV[CO_2_, spV[O_2_, and RQ, or means ± SE of per-individual Arrhenius parameters for *E*_a_ and ln *b*_0_. Different lowercase letters within a row indicate significant pairwise differences among overwintering treatments based on Tukey-adjusted post hoc comparisons (α = 0.05; letters row-specific and do not imply comparisons across rows). Test statistics and *P*-values at the end of each row report overall treatment effects for the corresponding model. Sample sizes (*n*) indicate the number of individual queens contributing to each analysis; *n* varies for Arrhenius parameters because they were only estimated for individuals with complete temperature series.

|  |  | 2°C<br>variable | 2°C | 3°C | 4°C | 4°C<br>variable | *n* | statistic | *P* |
| --- | --- | --- | --- | --- | --- | --- | --- | --- | --- |
| spV □ CO<sub>2</sub> | spV □ CO<sub>2</sub><br>(mL CO<sub>2</sub> g □<sup>-1</sup><br>h □<sup>-1</sup>) | 0.670 ±<br>0.096 **a** | 0.302 ±<br>0.043 **b** | 0.365 ±<br>0.052 **b** | 0.610 ±<br>0.088 **a** | 0.421 ±<br>0.060 **ab** | 24 | $\chi^2 = 30.96$ | **<0.001** |
|  | Thermal<br>sensitivity<br>(*E<sub>a</sub>*, eV) | 1.35 ±<br>0.090 **b** | 1.08 ±<br>0.051 **a** | 1.23 ±<br>0.068 **a** | 1.34 ±<br>0.084 **b** | 1.02 ±<br>0.102 **a** | 18-22 | *F* = 3.53 | **0.0099** |
|  | Baseline<br>metabolism<br>(ln *b*<sub>0</sub>) | 54.3 ±<br>3.85 **b** | 42.6 ±<br>2.14 **a** | 48.8 ±<br>2.80 **a** | 53.5 ±<br>3.54 **b** | 39.8 ±<br>4.30 **a** | 18-22 | *F* = 3.66 | **0.0081** |
| spV □ O<sub>2</sub> | spV □ O<sub>2</sub><br>(mL CO<sub>2</sub> g □<sup>-1</sup><br>h □<sup>-1</sup>) | 1.068 ±<br>0.134 **a** | 0.458 ±<br>0.057 **b** | 0.599 ±<br>0.075 **bc** | 0.903 ±<br>0.115 **ad** | 0.670 ±<br>0.084 **cd** | 24 | $\chi^2 = 49.5$ | **<0.001** |
|  | Thermal<br>sensitivity<br>(*E<sub>a</sub>*, eV) | 0.96 ±<br>0.068 **c** | 0.58 ±<br>0.051 **a** | 0.63 ±<br>0.054 **a** | 0.85 ±<br>0.060 **b** | 0.86 ±<br>0.081 **b** | 18-22 | *F* = 6.82 | **<0.001** |
|  | Baseline<br>metabolism<br>(ln *b*<sub>0</sub>) | 55.1 ±<br>2.70 **c** | 38.8 ±<br>2.70 **a** | 41.1 ±<br>2.68 **a** | 49.6 ±<br>2.74 **b** | 49.6 ±<br>2.82 **b** | 18-22 | *F* = 6.91 | **<0.001** |
| RQ | | 0.660 ±<br>0.103 | 0.727 ±<br>0.114 | 0.627 ±<br>0.098 | 0.707 ±<br>0.112 | 0.652 ±<br>0.102 | 24 | $\chi^2 = 1.252$ | 0.0869 |

### Mean winter temperature induces metabolic suppression under warming

Queens overwintered at colder constant mean temperatures (2 and 3°C) showed lower metabolic rates at warmer test temperatures than queens held at a constant 4°C. At 11°C, spV[CO_2_ and spV[O_2_ were significantly lower in the constant 2 and 3°C treatments relative to the constant 4°C treatment (Fig. 2A,B; Table 1), consistent with temperature-dependent metabolic suppression under colder winter conditions.

Differences among constant treatments were smaller at colder test temperatures (−5, 2, and 4°C; Tables S2), but the divergence became clear under warming. These results indicate that mean winter temperature alone shifts the baseline position and temperature dependence of metabolism, with colder means inducing deeper suppression.

### Thermal variability modifies mean-dependent suppression in opposite directions

The effects of thermal variability depended strongly on mean temperature. Queens overwintered under thermally variable conditions centered on 2°C showed substantially higher metabolic rates at 11°C than queens held at constant 2 and 3°C (Fig. 2A,B; Table 1; Supplemental Fig. 1). Thermal variability around 2°C therefore eliminated metabolic suppression observed at constant 2°C and resulted in metabolic rates comparable to those of queens overwintered at a constant 4°C.

Arrhenius analyses revealed that queens from the variable 2°C treatment showed elevated baseline metabolism and higher thermal sensitivity of mass-specific oxygen consumption relative to constant 2 and 3°C treatments (Fig. 3B,D; Table 1; Supplemental Fig. 2), indicating that variability at low mean temperature disrupts mean-induced metabolic suppression. Further, baseline rates and thermal sensitivity of mass-specific CO_2_ production were significantly higher for queens from the variable 2°C treatment relative to those from the variable 4°C treatment, suggesting positive metabolic compensation induced by variable 2°C conditions.

**Figure 3.**
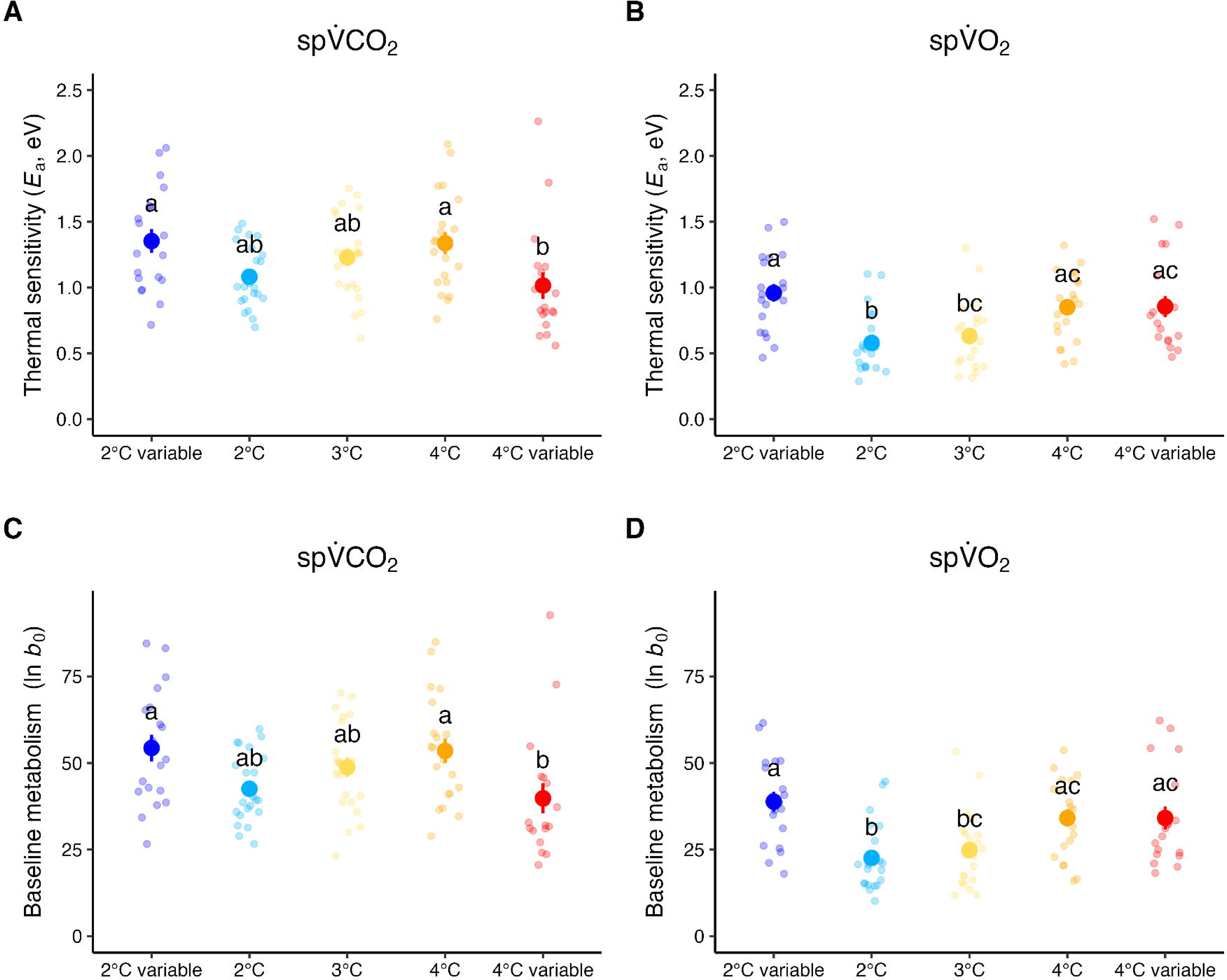
Overwintering thermal experience alters the thermal sensitivity of metabolism. The thermal sensitivity (*E*_a_, slope; top plots) and intercept (bottom plots) derived from Arrhenius fits for mass-specific CO_2_ production (A, C) and O_2_ consumption (B, D) in relation to temperature for queens overwintered in each of five thermal regimes. Small points are values for individual queens, and large points and error bars indicate group means ± 95% confidence intervals. Letters denote significant differences among regimes based on post hoc comparisons. See text and Table 1 for details.

In contrast, queens exposed to thermal variability centered on 4°C showed reduced baseline metabolic rates and lower thermal sensitivity relative to queens held at a constant 4°C (Fig. 3A,C; Table 1; Supplemental Fig. 2). This suppression was evident across much of the tested temperature range and was particularly pronounced in oxygen consumption at −5°C, where the variable 4°C treatment exhibited significantly lower spV[O_2_ than several other temperature treatments (Fig. S1B; Table S2). Together, these results suggest that variability at higher mean temperature enhances metabolic suppression rather than elevating energetic demand.

### Treatment effects are modest at cold test temperatures

At −5°C, most treatment differences were smaller and differed between gas-exchange metrics. Queens overwintered at constant 3°C showed lower spV[CO_2_ than queens from the variable 4°C treatment (Table S2), consistent with mean-dependent suppression. In contrast, the variable 4°C treatment showed reduced spV[O_2_ relative to other treatments, indicating metric-specific divergence under extreme cold (Fig. 2A,C; Table S2).

At intermediate test temperatures (2 and 4°C), treatment effects were smaller than those observed at 11°C, though significant differences persisted for spV[CO_2_ at both temperatures (Tables S2). At 4°C, spV[CO_2_ remained higher in the variable 2°C treatment relative to the variable 4°C treatment (Table S2), reinforcing the divergence between low-mean and high-mean variability responses.

### Minor variation in assay temperatures does not explain metabolic rate differences

Because acclimation and soak temperatures during respirometry deviated slightly from the nominal assay temperatures, we tested whether these deviations explained variation in metabolic rate (Supplemental Fig. 5). Models including acclimation temperature showed weak effects for CO_2_ production (*P* = 0.092) and a significant interaction between acclimation temperature and assay temperature for O_2_ consumption (*P* = 0.001). However, acclimation temperature alone was not significant for O_2_ consumption (*P* = 0.46). Models incorporating soak temperature yielded similarly weak results. Although soak temperature was marginally associated with CO_2_ production (*P* = 0.049), the interaction with assay temperature was not significant (*P* = 0.13), and no effects were detected for O_2_ consumption (all *P* > 0.09). Inspection of batch-specific relationships revealed no consistent directional effect (Supplemental Fig. 6), suggesting that minor variation in acclimation and soak temperatures did not systematically influence metabolic rate.

### Thermal variability alters the respiratory quotient (RQ) primarily in extreme cold

Respiratory quotient (RQ) varied strongly with measurement temperature and differed among overwintering treatments primarily at the coldest test temperature. At −5°C, queens overwintered under thermally variable conditions centered on 4°C had higher RQ values than queens from constant-temperature treatments (Fig. 2C; Table S2; Supplemental Fig. 1), indicating altered substrate use or gas exchange balance under extreme cold. In contrast, RQ did not differ appreciably among overwintering treatments at warmer test temperatures (Fig. 2C; Tables S2; Supplemental Fig. 1), suggesting that variability-dependent shifts in fuel use only occurred at the coldest temperatures.

### Thermal regime alters the proportional mass loss during diapause

Overwintering treatment significantly affected the proportion of body mass lost during diapause (quasibinomial GLM: *F*_4,115_ = 11.80, *P* < 0.0001; Fig. 4; Table S1; Supplemental Fig. 4). Post hoc comparisons revealed that queens from the variable 4°C treatment had significantly higher proportional mass loss than all other treatments except the constant 4°C treatment, and queens from the constant 4°C treatment lost proportionally more mass than those from the constant 3°C treatment (Fig. 4, Table S1). Proportional mass loss did not differ among the constant 2°C, variable 2°C, and constant 3°C treatments.

**Figure 4.**
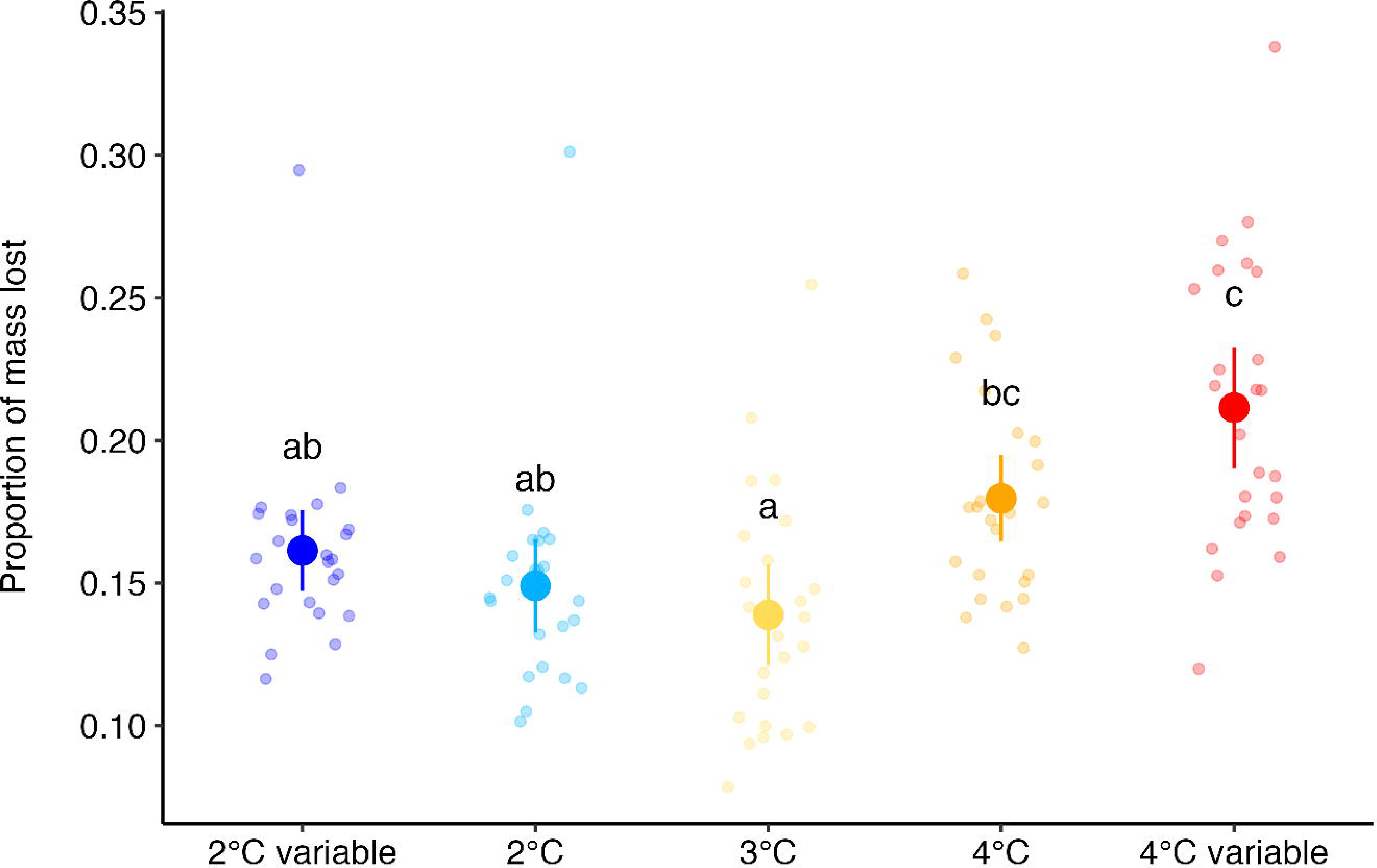
Proportional mass loss after six weeks differed among overwintering thermal regimes. Queens overwintered under variable temperatures centered on 4°C lost the most mass, whereas those at constant 2°C and 3°C lost the least. Those experiencing variable temperatures centered on 2°C did not differ in mass loss from those at constant 4°C. Transparent points show individual queens, and larger points and error bars show group means ± 95% confidence intervals. Letters indicate significant differences among regimes based on *post hoc* comparisons.

Pre-diapause mass did not significantly influence proportional mass loss, nor did its effect differ among treatments (all *P* > 0.35; Supplemental Fig. 4), indicating that treatment effects on mass loss were independent of initial body mass.

### Predicted survival consequences of altered metabolism

The divergent metabolic trajectories associated with overwintering thermal history translated into contrasting energetic outcomes during diapause. Using treatment-specific Arrhenius relationships (Fig. 3; Table 1; Supplemental Fig. 2) to project lipid depletion under alternative future thermal scenarios, we found that queens from the variable 2°C treatment were consistently predicted to exhaust lipid reserves faster than queens overwintered at constant 2°C (Fig. 5A). Elevated baseline metabolism (log_10_ *b*_0_) and higher thermal sensitivity (*E*_a_) in this group resulted in disproportionately high energetic expenditure during warmer periods.

**Figure 5.**
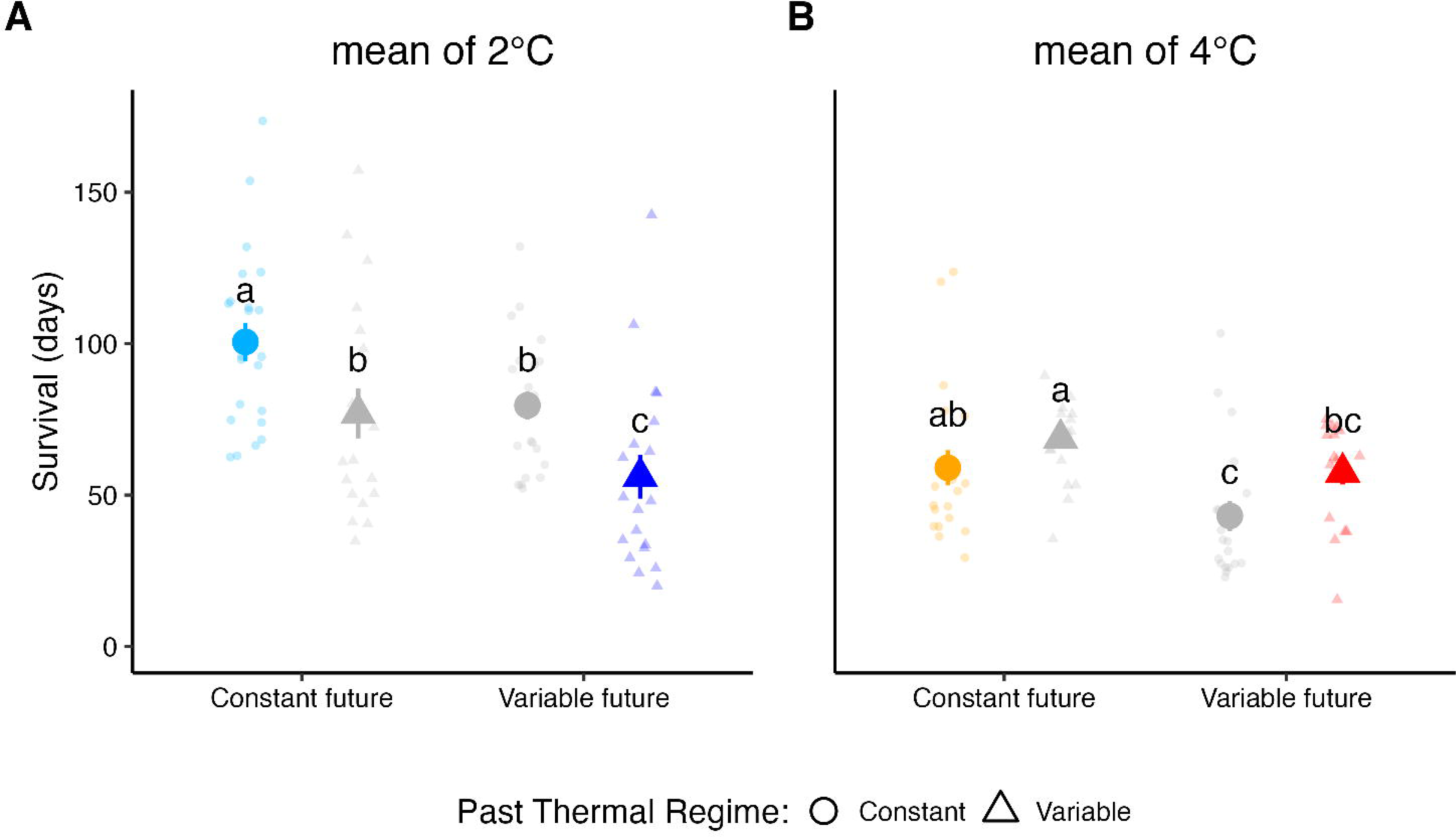
Temperature variability has context-dependent effects on overwinter survival. (A) For thermal regimes centered on 2°C, queens that experienced variable temperatures (± 6°C) during the first six weeks of diapause (triangles) are predicted to deplete lipid reserves sooner than those that experienced constant temperatures (circles) under both constant (gray triangle) and variable (blue triangle) future conditions. (B) Queens that experienced variable temperatures centered on 4°C (triangles) would survive similarly under constant and variable future conditions, whereas those exposed to constant thermal regimes would have significantly shorter survival times under variable relative to constant future conditions. Survival time was calculated as days to reach a critical lipid depletion threshold using Arrhenius-derived metabolic parameters (Fig. 3) and temperature trajectories representing constant or fluctuating future conditions (Fig. 1). Small points represent simulation outcomes for individual queens, and larger symbols indicate estimated marginal means ± 95% confidence intervals.

In contrast, queens exposed to variability around a mean of 4°C showed suppressed baseline metabolism and reduced thermal sensitivity relative to constant 4°C queens (Fig. 3A,C; Table 1; Supplemental Fig. 2). These shifts translated into lipid depletion rates and predicted survival durations that closely overlapped those of queens that had experienced a constant 4°C thermal regime (Fig. 5B).

Predicted survival time depended on both prior overwintering regime and subsequent thermal conditions (Fig. 5). For modeled scenarios with a mean of 2°C, queens that had experienced a variable thermal regime depleted lipid stores faster and died sooner than queens from constant regimes under both constant and variable future scenarios (Fig 5A, compare triangles to circles). Queens that had experienced a constant 2°C thermal regime were predicted to survive longest under constant future conditions, with shorter lifespans under variable future conditions.

Predicted survival patterns were completely different for modeled scenarios with a mean of 4°C (Fig. 5B). Queens that had experienced a variable 4°C thermal regime had no decrease in survival time under either constant or variable future conditions, suggesting that metabolic compensation in response to variable temperatures at this higher mean (Figs. 2, 3) can rescue queens from the energetic consequences of temperature variability (Fig. 1B).

Together, these results demonstrate that winter thermal variability does not uniformly accelerate energy use. Instead, its consequences depend on the mean-temperature context: variability centered on colder means can erode metabolic suppression and accelerate depletion, whereas variability centered on warmer means can enhance suppression and buffer energetic costs.

## Discussion

Winter temperature variability is a defining feature of natural environments, yet the energetic consequences during dormancy of this variability remain unclear. Because metabolic rate increases exponentially with temperature, variability in temperature will elevate cumulative energy expenditure (8–11, 21). But animals may compensate metabolically. Our results show that due to differential metabolic compensation, the energetic consequences of winter temperature variability depend on the mean temperature around which fluctuations occur.

Overwintering queens adjusted their metabolism in response to thermal experience, but the direction of this compensation depended on the mean temperature around which variability occurred: the same magnitude of fluctuation (± 6°C) produced contrasting energetic outcomes when centered on 2°C versus 4°C. Queens exposed to variability around 2°C exhibited elevated baseline metabolic rates and greater temperature sensitivity of metabolism when metabolic rate was measured across multiple assay temperatures (Figs. 2-3; Table 1). Under these conditions, variability increased metabolic rates during warming and accelerated predicted lipid depletion, with predicted downstream effects on overwinter survival (Fig. 5A). In contrast, queens exposed to variability centered on 4°C showed reduced baseline metabolism and lower thermal sensitivity relative to constant 4°C treatments (Fig. 3). This suppression corresponded to predicted survival trajectories that were similar to those of queens overwintered under constant 4°C conditions (Fig. 5B). Together, these results indicate that thermal variability can either amplify or reduce energetic demand depending on the mean temperature around which fluctuations occur.

During dormancy, the primary physiological challenge is conserving finite energy reserves that must sustain individuals through prolonged periods without feeding. Diapausing insects, therefore, typically suppress metabolism to minimize energetic expenditure and prolong survival (13, 16). In this context, physiological responses that reduce metabolic demand would be expected to buffer energetic costs during overwintering, rather than amplify metabolic expenditure to maintain physiological performance (27, 28). In our study, such suppression occurred under variability centered on 4°C, whereas variability centered on 2°C was associated with elevated metabolic demand.

Several physiological processes could contribute to the elevated metabolic demand observed under colder-centered variability. Repeated exposure to extreme low temperatures may impose physiological maintenance costs – such as maintaining ion homeostasis or repairing chill injury (25) – that could restrict metabolic suppression. Alternatively, repeated transitions across physiological thresholds may maintain metabolic flexibility in a manner similar to cold compensation observed in active ectotherms. Such responses are plausible because dormancy is not physiologically static. Insects can remodel tissues and reorganize metabolic pathways during diapause; for example, Colorado potato beetles reduce mitochondrial density in flight muscle during diapause to lower energetic demand (29). Endocrine pathways – including insulin signaling, adipokinetic hormone, and neuropeptide F – also regulate lipid mobilization and metabolic suppression during diapause (30–34). Although we did not examine these mechanisms directly, such processes provide plausible pathways through which thermal history could influence metabolic phenotype during dormancy.

Differences in metabolic parameters among overwintering treatments remained detectable when queens were later assayed across the range of test temperatures, suggesting that early thermal exposure *may* influence subsequent metabolic responses. Autumn temperatures are often more variable than those of mid-winter (20, 35), raising the possibility that conditions early in diapause could influence energetic trajectories later in the overwintering period. In our lipid depletion simulations, queens exposed to variability centered on 2°C consistently showed shorter predicted survival times across future thermal scenarios, whereas queens exposed to variability centered on 4°C exhibited survival trajectories similar to those of queens held under constant 4°C conditions (Fig. 5). However, whether these metabolic signatures persist throughout the entire overwintering period remains uncertain.

For bumble bee queens, overwinter survival determines whether colonies are established in spring. Climate impacts on bumble bees are often discussed in the context of summer heat stress (36, 37), yet winter temperature variability may represent an additional source of energetic constraint. If early overwinter conditions increase baseline metabolism or amplify metabolic responses to subsequent warming events, queens may enter late winter or early spring with reduced energetic reserves. As climate change alters both seasonal means and patterns of temperature variability, understanding how variability interacts with physiological processes may improve predictions of overwinter survival.

## Conclusion

The energetic consequences of winter temperature variability depend on the mean temperature around which fluctuations occur. In our study, variability centered on colder mean temperatures was associated with increased metabolic demand and shorter predicted survival, whereas variability centered on warmer mean temperatures corresponded with greater metabolic suppression and more stable survival trajectories. These findings suggest that evaluating the effects of climate variability on overwintering ectotherms requires considering not only the magnitude of temperature fluctuations but also the mean thermal context in which variability occurs.

## Materials and Methods

### Animals

We kept two commercial *Bombus impatiens* colonies (Natupol Excel, Koppert Biological Systems Inc., Howell, MI, USA) under controlled conditions in incubators (28°C, 12:12 light-dark cycle, 63% RH; I36VL, PERCIVAL Scientific, Inc., Perry, IA, USA) and supplied them with ground pollen (BZ Bodies, Clovis, CA, USA) and *ad libitum* access to Koppert’s supplied nectar.

### Experimental Setup

We marked callow bumble bee queens (easily identified by their silvery pile and curled wings) with color-numeric tags within two days of eclosion to track age and returned them to their natal colonies. After marking a sufficient number of queens (*n* = 120), we housed them in colony-specific boxes supplied with *ad libitum* nectar and returned them to incubators. We induced dormancy by gradually cooling incubators from 28°C to 4°C and under an 8:16 L:D cycle over seven days, followed by holding at 4°C in the dark for four more days. We then placed queens individually in 5-dram plastic vials (Thornton Plastics, Salt Lake City, UT, USA); each vial was connected via holes in the lid to a second 5-dram vial filled with dampened sphagnum moss to maintain humidity. Queens were haphazardly assigned to one of five temperature treatments, maintained in incubators: constant 2°C (2.01 ± 0.47°C), constant 3°C (3.0 ± 0.27°C), constant 4°C (4.02 ± 0.54°C), or two oscillating treatments. The first oscillating treatment ranged from −1 to 11°C (“4 ± 6°C”; 4.14 ± 5.74°C), while the second ranged from −5 to 7°C (“2 ± 6°C”; 1.77 ± 6.51°C; see Fig. 1C, Supplemental Figure 3).

### Metabolic Test Temperatures

After six weeks in dormancy, we measured the metabolic rate of each queen (*n* = 120; 24 per treatment) at each of four test temperatures (480 total metabolic measurements): −5, 2, 4, and 11°C, representing mean holding temperatures and the extreme values experienced in the oscillating treatments (Fig 1C). We monitored temperature during metabolic trials with T-type thermocouples connected to loggers recording every ten seconds (Onset HOBO 4-channel thermocouple data logger, Model: UX120-014M; Onset Computer Corporation, Bourne, MA, USA).

The 2, 4, and 11°C measurements were conducted in incubators (Percival Scientific, Perry, IA, USA), and the −5°C measurements were conducted in a chest freezer outfitted with a temperature controller (TIC-308, INKBIRD Tech. C.L., Luohu District, Shenzhen, Guangdong Province, China). We chilled the air used to flush test chambers by first flowing it through copper coils (internal diameter = 7.82 mm, outer diameter = 9.6 mm, length = 1.25 m) submerged in large containers of windshield fluid (3.5 L per container, non-freezing) kept in each chamber. This chilled the air appropriately before it reached a homerun manifold through which we flushed 12 syringes simultaneously; 10 containing queens and two left empty to serve as calibration measurements before and after all 10 queens in a trial.

### Respirometry

Each 30-mL syringe had a small hole drilled near the plunger opening and a two-way valve attached to the tip. By opening all two-way valves connected to the manifold, and drawing the plungers out past the drilled holes, we flushed all syringes for two minutes with dry, CO_2_-free air (2 L min^-1^) from an FT-IR Purge Gas Generator (Parker-Hannifin Corporation, Cleveland, OH, USA) conditioned at chamber temperature.

After flushing, we depressed each plunger past the drilled hole, closed each valve to seal the syringe, and noted the initial air volume. We conducted preliminary trials to determine the maximum duration we could seal bees without inducing hypoxia or hypercapnia, while still ensuring a measurable respiratory signal. For example, at 2°C, syringes sealed for two hours had oxygen depletion less than 2% and CO_2_ accumulation below 1600 ppm. At 11°C, we limited sealing time to 60 minutes to avoid exceeding these thresholds. Based on these trials, we sealed syringes for 90 minutes at −5, 2, and 4°C, and for 60 minutes at 11°C.

We measured CO_2_ production and O_2_ consumption using stop-flow respirometry with an integrated respirometry system (FMS-3; Sable Systems International Inc., Las Vegas, Nevada, USA). We zeroed (using pure N_2_ gas) and spanned the CO_2_ and O_2_ sensors daily with a primary standard (1011.11 ppm) and outside air (assumed value: 20.95%), respectively. For measurements of syringe gases, we flowed nitrogen (N_2_) from a gas cylinder through a column of DRIERITE–Ascarite II–DRIERITE to scrub residual water vapor and CO_2_. Flow rate was regulated at 150 mL min^-1^ by the FMS-3’s built-in mass-flow controller before the dry, CO_2_-free air passed sequentially through the flow meter, CO_2_ sensor (fractional concentration), and O_2_ sensor (%). Air samples were injected into a three-way valve located between the flow meter and the WVP sensor. To prevent dilution effects from residual moisture or CO_2_ from each syringe injection (Lighton, 2018), we plumbed a 30-mL column of magnesium perchlorate (Mg(ClO_4_)_2_) before the CO_2_ sensors to remove water vapor, followed by a 30-mL column of Ascarite II–Mg(ClO_4_)_2_ between the CO_2_ and O_2_ sensors to scrub CO_2_.

We first injected an empty chamber sealed just prior to the first chamber holding an animal, followed by sequentially injecting all 10 chambers containing animals, followed by an empty chamber sealed immediately after the last animal chamber. For each injection, we slowly injected 10 mL of syringe air into the N_2_ stream via the three-way valve.

### Respirometry analyses

We determined the syringe seal time as the fractional hours from sealing to injection (± 2 seconds). We adjusted the initial and final volumes to account for the body volume of each queen (estimated from bee mass measured immediately before respirometry and an assumed tissue density of 1 g cm^-3^) and defined injection periods as the time from the start of the injection to the opening of the three-way valve for the next injection, with the final peak ending 2.5 minutes after injection (the time-constant for washout given the volume of the respirometry setup and flow rate was approximately 2 minutes). We corrected the baseline drift of both O_2_ and CO_2_ sensors by adjusting the trace based on the slope between the gas concentrations before and after each peak. We converted the drift-corrected values to fractional proportions, multiplied by flow rates (mL min^-1^), divided by 60 to convert to mL sec^-1^ for second-by-second readings, and summed to estimate total gas volumes in the injected sample. We calculated the Respiratory Quotient (RQ) as the ratio of mass-specific CO_2_ production to O_2_ consumption (spV[CO_2_ and spV[O_2_, respectively).

Using each queen’s thermal response curves for mass-specific gas exchange, we calculated activation energy (*E*_a_), which describes how sensitive metabolic rate is to temperature changes, and the intercept (log *b*_0_), which represents the baseline metabolic rate at a standard reference temperature, to compare across overwintering thermal regimes, and across taxa (38, 39).

### Overwintering lipid depletion simulations

To predict ecological consequences of temperature-dependent metabolic responses, we simulated lipid depletion in bumble bee queens using empirically derived metabolic parameters under alternative overwintering scenarios. After six weeks of exposure to the overwintering treatments, we quantified metabolic responses and used those parameters to simulate subsequent lipid depletion and energetic consequences.

We modeled temperature-dependent CO_2_ production for each queen using Arrhenius relationships fit to mass-specific spV[CO_2_ measurements (Fig. 2; Supplemental Fig. 2). For each individual, we used the Arrhenius intercept (*b*_0_) and slope to predict whole-organism CO_2_ production across hourly temperature profiles. We calculated spV[CO_2_ (assuming body mass of 0.5 g, near the mean body mass for queens in this experiment) at hourly resolution and summed these values to estimate daily CO_2_ production. We then converted daily CO_2_ production to lipid consumption using an energetic equivalent of 27.9 J mL^-1^ CO_2_ and an energy density of 37 kJ g^-1^ of lipid (Lighton, 2018).

We ran simulations under four future thermal scenarios: constant 2°C and 4°C, and variable 2°C and 4°C (± 6°C; hourly temperatures with a sinusoidal cycle). For each combination of experienced thermal regime during the initial six-week exposure and future thermal scenario, we used the cumulative sum of daily lipid depletion, assuming bees started lipid stores equal to 10% of body mass to determine when lipid stores would fall under to 2% of body mass, a threshold associated with diapause termination and forced emergence (20, 40). We defined the number of days required to reach this threshold as the predicted survival time. To evaluate the influence of thermal history on subsequent energetic outcomes, we predicted survival times under both a continuation of previous thermal regimes (e.g., queens exposed to the variable 2°C treatment continuing to experience variable 2°C conditions) and under switched thermal regimes (e.g., bees that had experienced 2°C variable conditions subsequently experiencing constant 2°C).

### Statistical Analyses

We analyzed the data in R (version 4.4.3) (41) with packages *tidyverse* (42), *glmmTMB* (43), *lme4* (44), *lmerTest* (45), *car* (46)*, emmeans* (47), and supporting packages for visualization and post hoc inference. We conducted all hypothesis tests using a two-tailed significance level of α = 0.05.

### Body mass and proportional mass loss

To quantify changes in body mass during the six-week overwintering exposure, we summarized each queen’s mass as: (1) pre-diapause mass, and (2) post-treatment mass, defined as the mass measured immediately before the first respirometry trial. For each queen, we calculated absolute mass loss as the difference between pre-diapause mass and post-treatment mass (Δmass = pre − post), and proportional mass loss as absolute mass loss standardized by pre-diapause mass (Δmass / pre).

We tested whether pre-diapause mass (recorded before dormancy induction) and post-diapause (measured after 6 weeks of dormancy, immediately before the first respirometry trial) differed among overwintering treatments using linear models, followed by Tukey-adjusted pairwise comparisons using estimated marginal means, where appropriate. We tested whether pre-diapause mass predicted proportional mass loss (pre - post / pre) and whether this relationship differed among treatments using a generalized linear model (GLM) with a quasi-binomial error distribution, and compared slopes among treatments using *emtrends* (emmeans) on the fitted interaction model. For post-hoc contrasts of quasi-likelihood models, we report unadjusted pairwise comparisons because quasi families lack a true likelihood and Tukey-style multiplicity adjustments are derived under full likelihood assumptions (47, 48).

### Chamber temperature covariate during respirometry trials

To evaluate whether small deviations in chamber temperature during trials influenced metabolic measurements, we quantified two temperature covariates for each queen and nominal test temperature: (1) the mean “acclimation” temperature experienced between placement in the syringe and sealing after flushing, and (2) the mean “soak” temperature experienced during the sealed interval before gas injection. We tested whether variation in these temperatures explained residual variation in mass-specific CO_2_ production (spV[CO_2_) and O_2_ consumption (spV[O_2_) using generalized linear mixed-effects models with gamma-error distributions. These models included individual identity and experimental batch as random effects to account for repeated measurements on individuals and potential run effects.

### Metabolic rate responses to overwinter thermal regime and test temperature

We analyzed mass-specific CO_2_ production (spV[CO_2_) and O_2_ consumption (spV[O_2_) using generalized linear mixed-effects (GLME) models with gamma-error distributions. Fixed effects included overwintering treatment, test temperature, and their interaction to assess whether thermal history altered both the magnitude and temperature dependence of metabolic rate. We included individual identity as a random effect to account for repeated measurements of each queen across metabolic test temperatures and included batch and run (nested within batch) as random effects to account for experimental structure.

We evaluated the significance of fixed effects using Type III Wald χ^2^ tests. When significant interactions were detected, we conducted post-hoc comparisons among overwintering treatments within each test temperature using estimated marginal means with Tukey-adjusted multiple-comparison procedures.

### Respiratory Quotient (RQ; CO_2_:O_2_)

We analyzed RQ using generalized linear models with a gamma-error distribution, with overwintering treatment, test temperature, and their interaction as predictors. We evaluated model effects using Type III tests, and when interactions were significant, we conducted treatment contrasts within each test temperature using estimated marginal means with Tukey-adjusted multiple-comparison procedures.

## Acknowledgements

We thank M. K. Dillon for insightful and constructive feedback on the experimental design and its justification, and C. M. F. Glass for editorial comments that improved the manuscript. We are also grateful to T. R. Kircher, J. O. Ray, C. M. Smith, D. J. Stevens, and M. A. Stock for their support during the analysis and writing of this manuscript. A. T. Cressman provided valuable assistance with experimental preparation and the handling and tagging of callow queens. This work was supported by the National Science Foundation (grant numbers EF-1921562, OIS-1826834 to M.E.D.) and by the University of Wyoming Research and Economic Development Division (Teton Postdoctoral Fellowship to J.R.G.).

