## Supplemental Materials for "Mean temperature determines whether winter variability accelerates or buffers energy loss"

**Supplemental Tables**

**Supplemental Table 1.** Overwintering-associated mass loss in *Bombus impatiens* queens across thermal treatments. Values are means ± SE. Change in mass (Δ mass) represents absolute mass loss during overwintering, and the proportion lost is Δ mass divided by pre-overwintering mass. Different lowercase letters indicate significant pairwise differences among treatments. Post mass and Δ mass were analyzed using one-way ANOVA with post hoc tests, while proportion lost was analyzed using a quasibinomial model; estimates are shown on the response scale (for this model, post-hoc contrasts reflect overall clustering as some comparisons are not transitive). The *F* and P values indicate overall treatment effects.

|  | 2°C | 2°C variable | 3°C | 4°C | 4°C variable | *F* | *P* |
| --- | --- | --- | --- | --- | --- | --- | --- |
| ***n*** | 24 | 24 | 24 | 24 | 24 | — | — |
| **Pre mass (g)** | 0.578 ± 0.013 | 0.560 ±  0.015 | 0.593 ±  0.013 | 0.601 ±  0.014 | 0.563 ±  0.011 | 1.87 | 0.121 |
| **Post mass (g)** | 0.485 ± 0.013 **ab** | 0.477 ±  0.014 **ab** | 0.510 ±  0.012 **b** | 0.493 ±  0.013 **b** | 0.444 ±  0.009 **a** | 4.06 | **0.004** |
| **Δ mass (g)** | 0.083 ± 0.004 **ab** | 0.093 ±  0.004 **a** | 0.082 ±  0.006 **a** | 0.108 ±  0.005 **bc** | 0.120 ±  0.007 **c** | 9.79 | **<0.001** |
| **Proportion lost** | 0.149 ± 0.008 **ab** | 0.161 ±  0.007 **ab** | 0.139 ±  0.009 **a** | 0.180 ±  0.007 **bc** | 0.211 ±  0.010 **c** | 11.8 | **<0.001** |

**Supplemental Table 2.** Mass-specific rates of CO_2_ production (spV̇CO_2_), O_2_ consumption (spV̇O_2_), and respiratory quotient (RQ) measured at −5 °C, 2 °C, and 4 °C across overwintering treatments. Values are estimated marginal means ± SE. Test statistics and *P*-values (α = 0.05) report overall treatment effects for each response variable at each test temperature. Different lowercase letters within a row indicate significant pairwise differences among treatments based on Tukey-adjusted post hoc comparisons (α = 0.05). Rows without letters indicate no significant pairwise differences.

|  |  | **2°C variable** | **2°C** | **3°C** | **4°C** | **4°C variable** | Test | P |
| --- | --- | --- | --- | --- | --- | --- | --- | --- |
| Response @ -5°C | **spV̇CO**_2_ | 0.0278 ± 0.0041 **ab** | 0.0224 ± 0.0032 **ab** | 0.0187 ± 0.0027 **a** | 0.0231 ± 0.0033 **ab** | 0.0364 ± 0.0055 **b** | *χ*^2^ = 15.88 | **0.003** |
|  | **spV̇O**_2_ | 0.1177 ± 0.0150 **a** | 0.1164 ± 0.0145 **a** | 0.1191 ± 0.0148 **a** | 0.1037 ± 0.0129 **ab** | 0.0771 ± 0.0102 **b** | *χ*^2^ = 13.03 | **0.011** |
|  | **RQ** | 0.281 ± 0.0457**a** | 0.214 ± 0.0338 **ab** | 0.169 ± 0.0267 **b** | 0.240 ± 0.0380 **ab** | 0.577 ± 0.0959 **c** | *χ*^2^ = 62.39 | **< 0.0001** |
| Response @ 2°C | **spV̇CO**_2_ | 0.0393 ± 0.00589 | 0.0366 ± 0.00525 | 0.0548 ± 0.00795 | 0.0482 ± 0.00718 | 0.0345 ± 0.00503 | *χ*^2^ = 9.52 | **0.0493** |
|  | **spV̇O**_2_ | 0.153 ± 0.0201 | 0.129 ± 0.0164 | 0.139 ± 0.0178 | 0.146 ± 0.0193 | 0.137 ± 0.0176 | *χ*^2^ = 1.736 | 0.784 |
|  | **RQ** | 0.317 ± 0.0524 | 0.337 ± 0.0539 | 0.471 ± 0.0756 | 0.378 ± 0.0617 | 0.363 ± 0.0607 | χ² = 6.92 | 0.1401 |
| Response @ 4°C | **spV̇CO**_2_ | 0.070 ± 0.010 **a** | 0.052 ± 0.007 **ab** | 0.059 ± 0.008 **ab** | 0.051 ± 0.007 **ab** | 0.038 ± 0.005 **b** | *χ*^2^ = 13.68 | **0.008** |
|  | **spV̇O**_2_ | 0.211 ± 0.027 | 0.147 ± 0.019 | 0.167 ± 0.021 | 0.147 ± 0.018 | 0.145 ± 0.018 | *χ*^2^ = 10.74 | **0.030** |
|  | **RQ** | 0.346 ± 0.0545 | 0.394 ± 0.0626 | 0.382 ± 0.0608 | 0.370 ± 0.0582 | 0.298 ± 0.0476 | χ² = 3.96 | 0.412 |

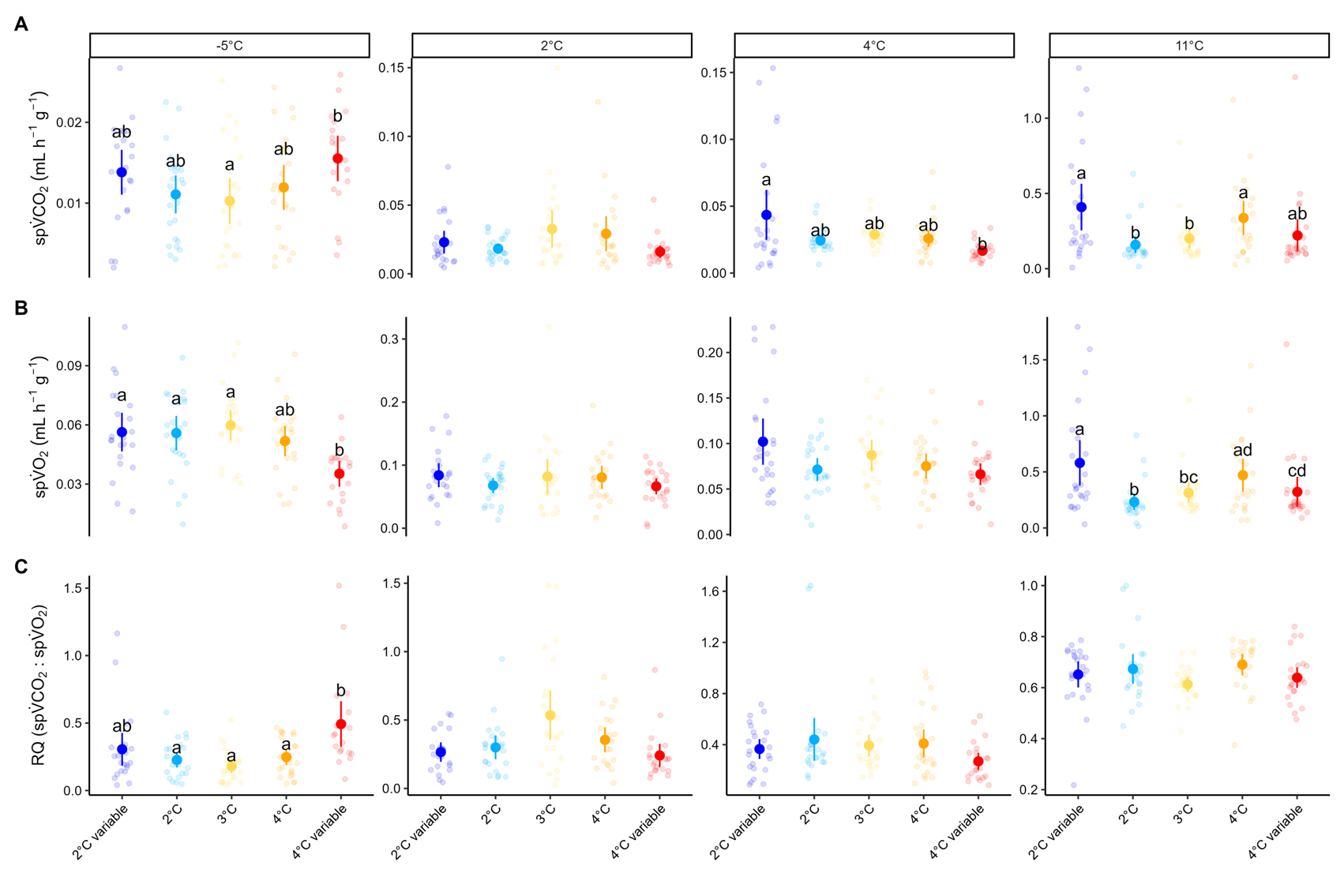
**Supplemental Figures**

**Supplemental Figure 1**. Overwintering treatment effects on mass-specific metabolic rates and respiratory quotient across test temperature. Mass-specific rates of (**A**) CO_2_ production (spV̇CO_2_) and (**B**) O_2_ consumption (spV̇O_2_), and (**C**) respiratory quotient (RQ = spV̇CO_2_ : spV̇O_2_) are shown for queens from each overwintering treatment at four test temperatures (-5, 2, 4, and 11°C). Points represent individual measurements (jittered). Large circles and error bars show treatment means ± 95% CI. Different letters denote significant pairwise differences among overwintering treatments within a given test temperature based on Tukey-adjusted post hoc comparisons (*α* = 0.05). RQ values greater than 1.8 are included in our analyses, but are not shown for ease of comparison.

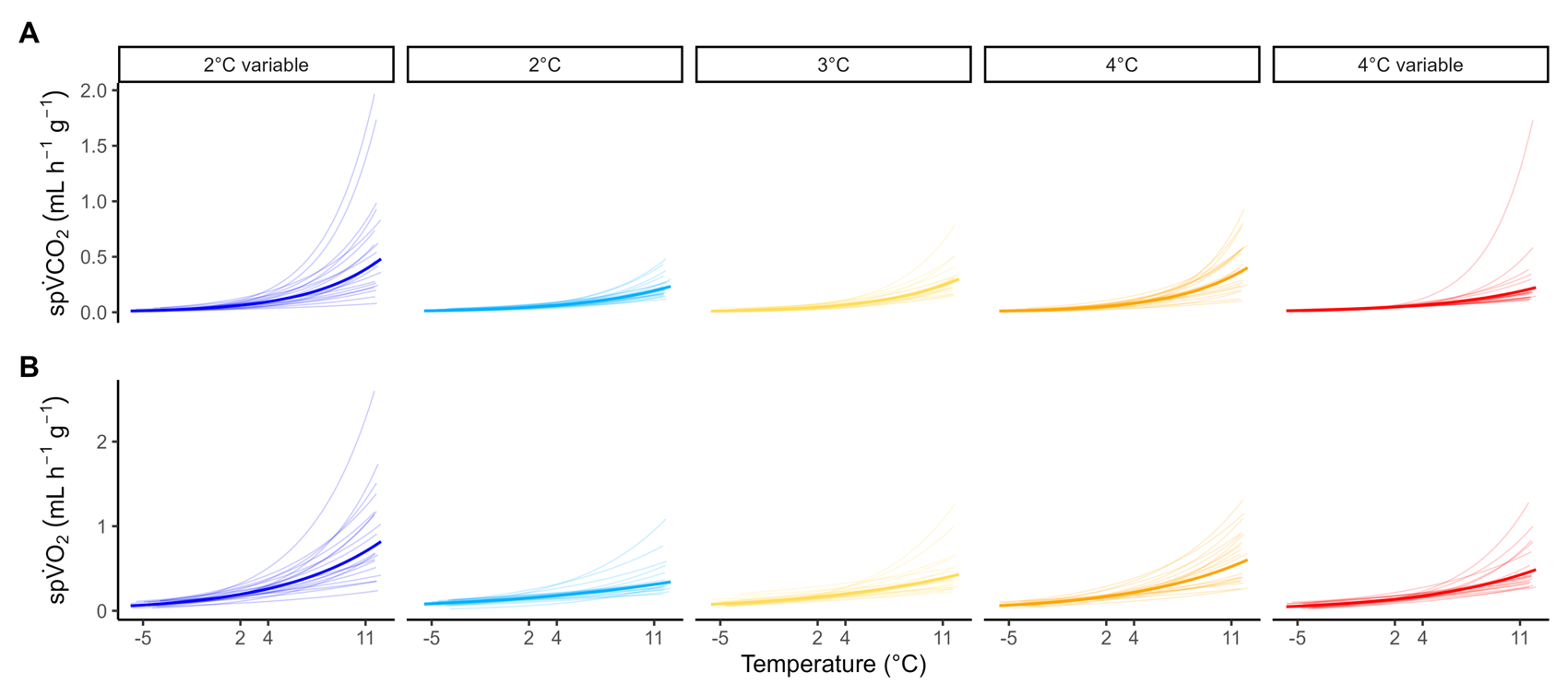
**Supplemental Figure 2**. **Treatment-level Arrhenius predictions with individual-level variation.** (**A**) Predicted spV̇CO_2_ as a function of test temperature. (**B**) Predicted spV̇O_2_ as a function of test temperature. Bold lines show treatment-level predictions; faint lines show per-bee Arrhenius predictions generated from individual fits across each bee’s observed test-temperature range.

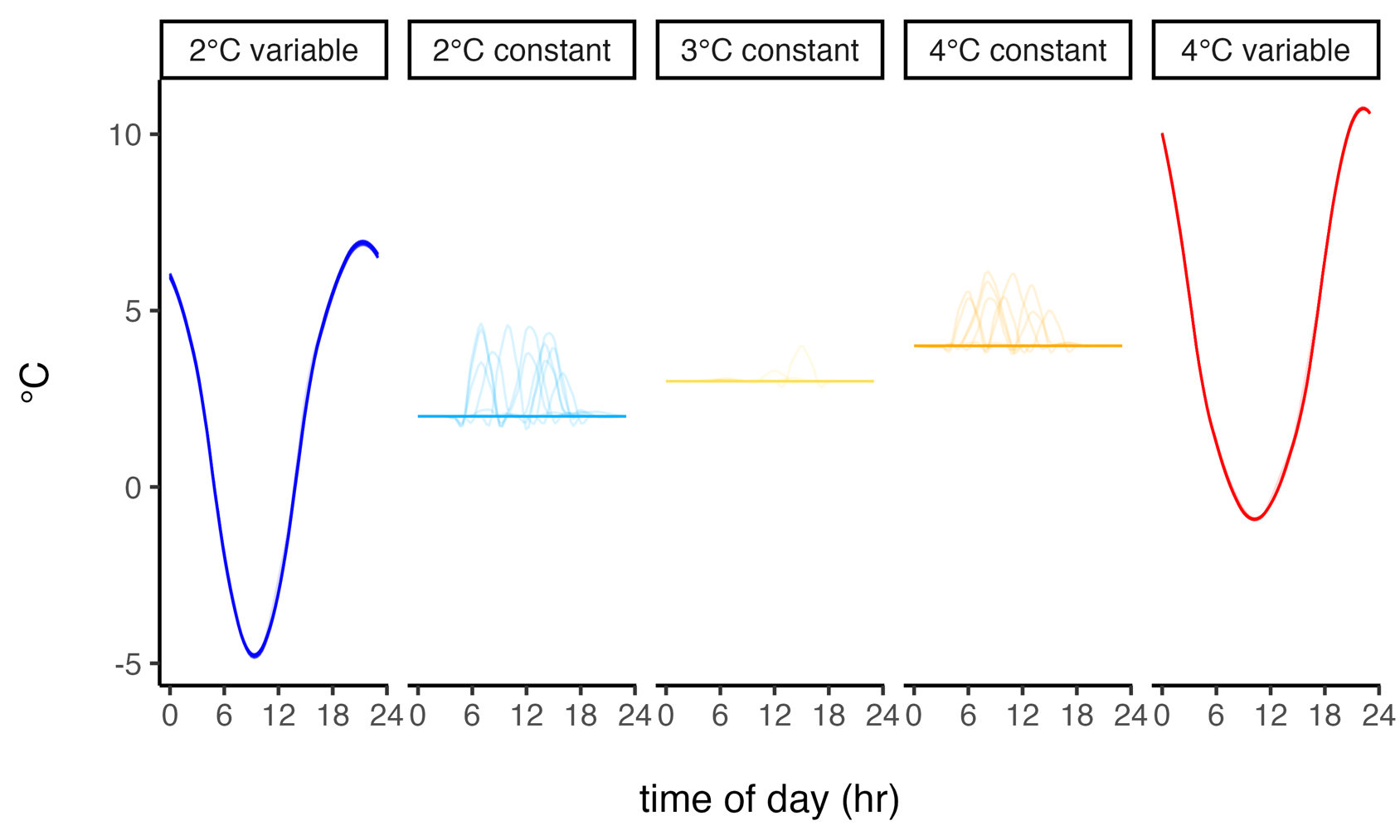
**Supplemental Figure 3**. Daily temperature profiles for overwintering treatments. Raw temperature recordings from overwintering temperature chambers were summarized to hourly mean temperature for each day and plotted across a 24-h cycle. Each line represents a single day. Panels correspond to the five overwintering treatments: 2°C variable, 2°C constant, 3°C constant, 4°C constant, and 4°C variable. Variable treatments fluctuated ± 6°C around their mean setpoints over a daily cycle, whereas constant treatments remained near their programmed temperatures, with deviations from the constant mean due to opening the door of the chamber during experimental runs.

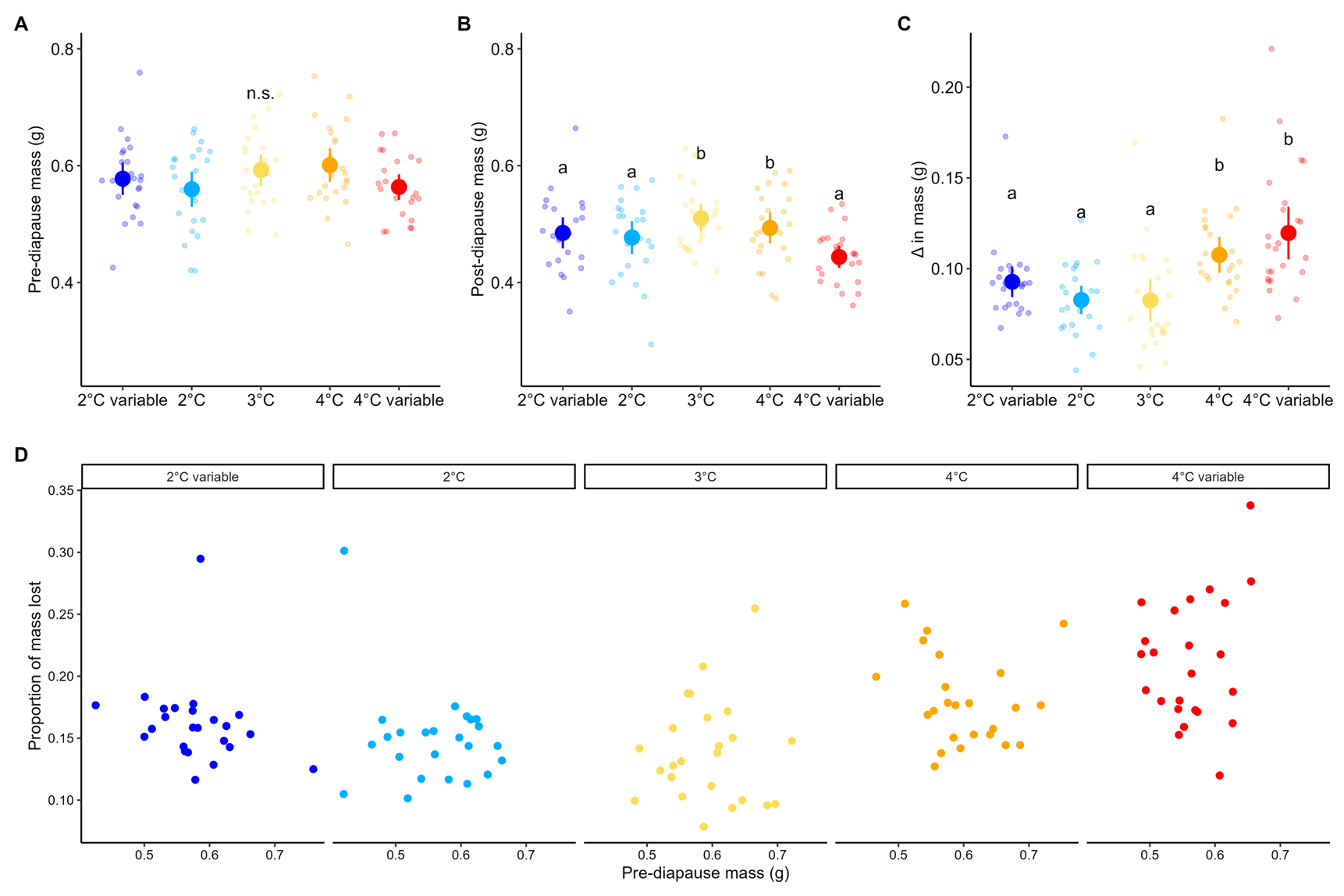

**Supplemental Figure 4**. Overwintering thermal regime influences mass loss but not initial mass. (**A**) Pre-diapause mass (body mass at the beginning of the experiment) did not differ among treatments. (**B**) Post-diapause mass (body mass after six weeks of exposure to the assigned overwintering regime and measured immediately before metabolic assays) differed among treatments. (**C**) Absolute mass loss during diapause (Δ mass = pre-diapause mass − post-diapause mass) varied among treatments. (**D**) Proportional mass loss (Δ mass / pre-diapause mass) plotted against initial mass for each treatment; no significant relationships with initial mass were detected. Points represent individual queens. Large circles and error bars indicate treatment means ± 95% CI. Different letters denote significant pairwise differences based on Tukey-adjusted post hoc comparisons (α = 0.05).

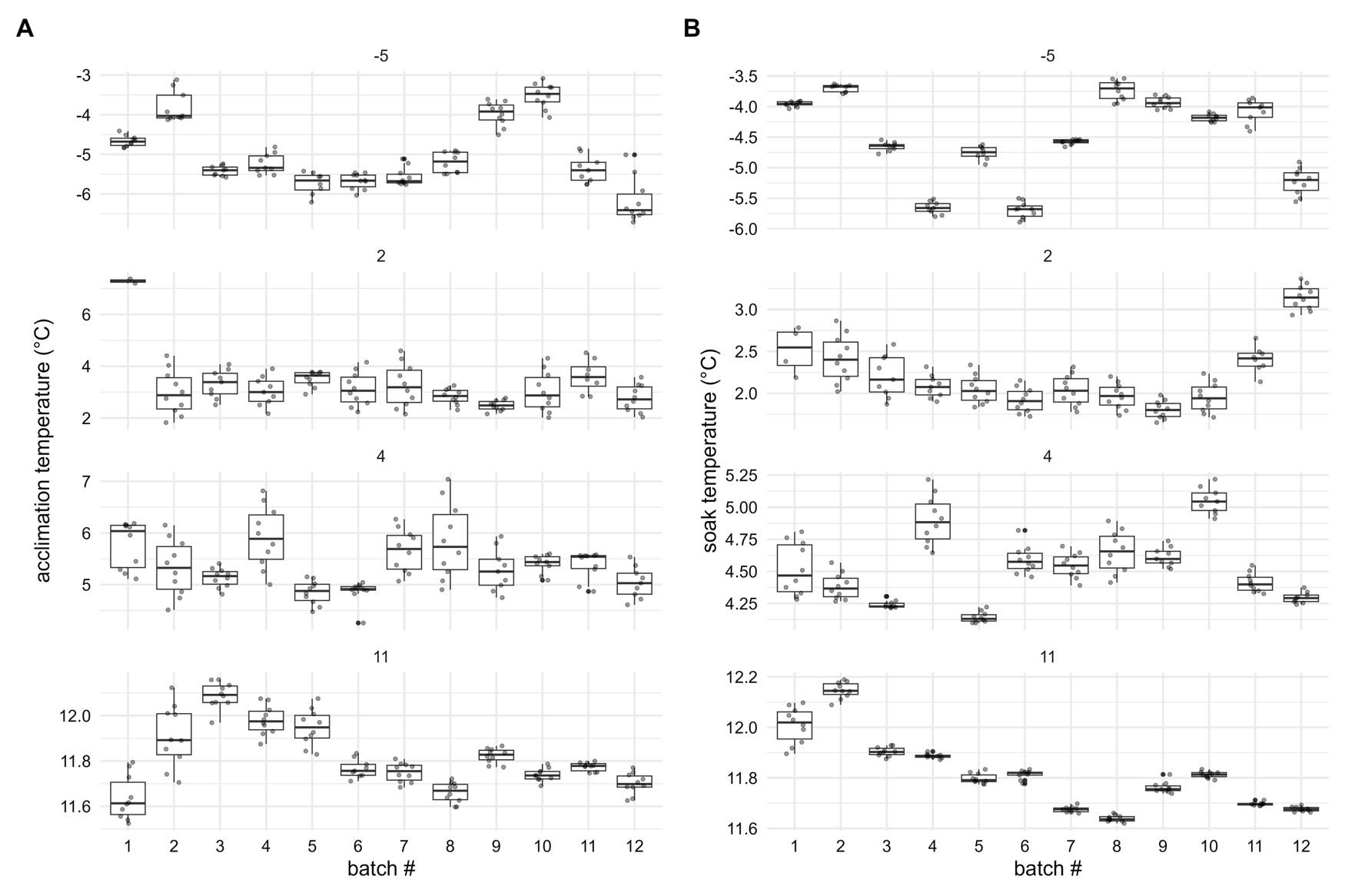

**Supplemental Figure 5**. To evaluate whether small temperature deviations during trials influenced metabolic measurements, we visualized (**A**) the mean acclimation temperature and (B) the mean soak temperature for each queen, grouped by experimental batch and faceted by test temperature (-5, 2, 4, 11°C). Acclimation temperature (A) represents the mean temperature of the air used to flush each syringe immediately before sealing (i.e., before the start of the metabolic measurements). Flushed air was equilibrated to chamber temperature by passing it through copper coils submerged in window-washing fluid, ensuring thermal matching with the set test temperature. Soak temperature (B) represents the mean incubator temperature during the sealed interval when queen gas exchange occurred inside the syringe. Boxplots show medians and interquartile ranges; points represent individual observations.

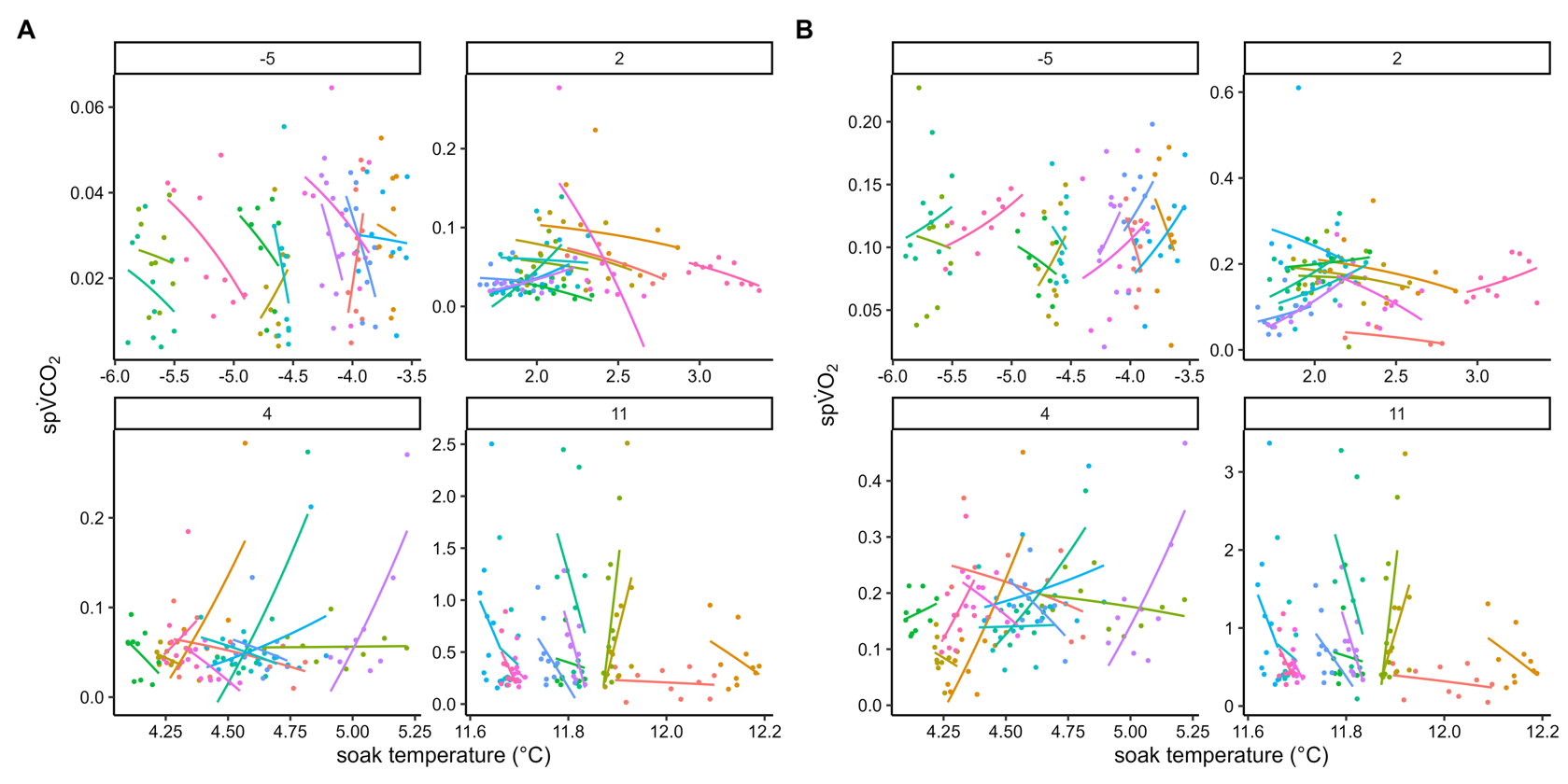

**Supplemental Figure 6**. Small variation in soak temperature did not systematically influence metabolic rate. To determine whether minor deviations in chamber temperature during trials affected metabolic measurements, we examined the relationship between mean soak temperature and mass-specific metabolic rate at each test temperature (-5, 2, 4, 11°C). (**A**) Mass-specific CO_2_ production (spV̇CO_2_) and (**B**) mass-specific O_2_ consumption (spV̇O_2_) plotted as a function of soak temperature. Soak temperature represents the mean incubator temperature during the sealed interval when gas exchange occurred inside the syringe. Points represent individual measurements and are colored by experimental batch. Panels are faceted by test temperature. Lines show fitted exponential relationships for visualization only.
